# Dorsal anterior cingulate cortex neurons only partially integrate determinants of value

**DOI:** 10.1101/2020.01.01.892380

**Authors:** Habiba Azab, Benjamin Hayden

## Abstract

Evaluation often involves integrating multiple determinants of value, such as the different possible outcomes in risky choice. A brain region can be placed either before or after a presumed evaluation stage by measuring how responses of its neurons depend on multiple determinants of value. A brain region could also, in principle, show partial integration, which would indicate that it occupies a middle position between (pre-evaluative) non-integration and (post-evaluative) full integration. Existing mathematical techniques cannot distinguish full from partial integration and therefore risk misidentifying regional function. Here we use a new Bayesian regression-based approach to analyze responses of neurons in dorsal anterior cingulate cortex (dACC) to risky offers. We find that dACC neurons only partially integrate across outcome dimensions, indicating that dACC cannot be assigned to either a pre- or post-evaluative position. Neurons in dACC also show putative signatures of value comparison, thereby demonstrating that comparison does not require complete evaluation before proceeding.

## INTRODUCTION

To evaluate an option, we must consider all of the aspects of the option that influence value and then combine them to generate an integrated value signal (Rangel et al., 2008; Kable & Glimcher, 2009; Busemeyer et al., 2019; Tversky, 1969; Tversky & Kahneman, 1974). For example, a risky prospect (a gamble) may involve two possible outcomes, both of which have some influence on our likelihood of choosing the gamble. To evaluate the gamble, then, we must decide how appealing each of those outcomes is and then somehow combine those component evaluations. This combination stage could involve a process of multiplication and addition, as dictated by normative approaches, or could be any of a number of satisficing heuristic approaches (Stewart et al., 2006; Busemeyer and Townsend, 1998; Tversky and Kahneman, 1974; Farashahi et al., 2019). Clearly, this integration process must be reified in the brain—we necessarily process dimensions separately and our actions reflect integrated values, so combination presumably has to occur somewhere in the middle (Bowman et al., 2012; Roesch and Olson, 2003; Roesch and Olson, 2004). How the integration occurs remains an important problem in neuroeconomics.

Indeed, neuroeconomists often seek to categorize brain regions as either preceding or following the integration process (Rangel et al., 2009). A standard approach is to probe the responses of single neurons and test whether their responses are driven by multiple determinants of value (see, e.g., Padoa-Schioppa and Assad, 2006; Kahnt et al., 2011; Hosokawa et al., 2013; Kennerley et al. 2009; Blanchard and Hayden, 2014; So and Stuphorn, 2010; O’Neill & Schultz, 2018; Raghuraman & Padoa-Schioppa, 2014). Simultaneous selectivity for multiple factors that determine value/utility has become a hallmark of cross-dimensional integration (Padoa-Schioppa, 2011; Hunt et al., 2015; Fellows, 2006; Kim et al., 2008; O’Neill and Schultz, 2018; Strait et al., 2014). However, this interpretation relies on the assumption that the process of evaluation is anatomically localized. Another possibility would be that it unfolds gradually across many regions, much as complex form is gradually built up over multiple regions of the ventral visual stream (Yoo and Hayden, 2018). That is, reward-sensitive regions may be have a hierarchical relationship, and the dimensions that influence value may become gradually more united the further we travel along it Given this possibility, it is notable that the standard statistical approach of detecting dependence on multiple determinants of value does not distinguish full (i.e. complete) from partial integration. We recently developed a method that makes use of Bayesian regression and cross-validation that can distinguish partial from complete correlations across dimensions in neural ensemble regressions, although we used it for a different purpose (Azab and Hayden, 2017). Here we use that methods to disambiguate partial from complete cross-dimensional integration.

A secondary benefit of our approach is that it allows us to address a question about how evaluation relates to comparison. Conventional approaches choice assumes the completion of evaluation before the start of comparison (Rangel et al., 2008, Padoa-Schioppa, 2011). For example, competitive inhibition models, the dominant models of economic choice, involve separate pools of integrated (i.e. post-evaluation) value neurons competing (Chau et al., 2014; Rustichini and Padoa-Schioppa, 2015; Hunt et al., 2012; Hunt et al., 2015; Louie et al., 2011). However, there is no special reason why comparison must wait until evaluation is complete (Yoo and Hayden, 2018). In distributed choice systems, such as beehives, comparison processes can progress even in the absence of full cross-dimensional integration. For example, individual bees that participate in a swarm decision may sample only a subset of available determinant dimensions and nonetheless contribution directly to comparison (Seeley, 2009; Seeley & Buhrman, 1999). It is not clear, in the human brain, whether evaluation is complete before comparison can begin, or whether comparison starts soon after evaluation does (Hunt et al., 2014; Cisek, 2012; Chen & Stuphorn, 2015; Strait et al., 2015; Azab & Hayden, 2018; Balasubramani et al., 2018). Cognitive models of value-based decision-making have, in fact, often proposed that the evaluation and comparison stages occur in parallel (Busemeyer et al., 2019; Noguchi & Stewart, 2018). Knowing whether signatures of integration across dimensions is partial or complete can help shed light on this question.

These questions are especially interesting to ask in the dorsal anterior cingulate cortex (dACC). Among value-sensitive or economic regions, there is a broad consensus that this is an anatomically or hierarchically late region (e.g. Heilbronner and Hayden, 2016; Rushworth et al., 2011; Schall et al., 2002; Wunderlich et al., 2009). That is, it may serve as a repository for feed-forward signals generated by other value-sensitive regions, such as orbitofrontal cortex and ventromedial prefrontal cortex, and have a direct role in influencing motor and premotor processing that other economic regions lack (Hayden et al., 2010). As a consequence, it stands to reason that, among value-sensitive regions, it should be most likely to carry fully-integrated signals. Indeed, current debates about the economic function of the dACC are generally split between (late) post-comparison selection role and (middle) post-evaluative comparison role (Hare et al. 2011; Wunderlich, Rangel & O’Doherty, 2009; Rangel & Hare, 2010; Azab & Hayden, 2017).

Here, we analyzed responses of neurons in dACC during a two-option token gambling task (Azab & Hayden, 2017; Azab & Hayden, 2018). We find that while neurons encode both large and small prospective outcomes of each gamble, the manner in which they encode these outcomes (the population vectors of their tuning slopes) are more similar than would be expected by chance if they were unrelated. Simultaneously, they are more *dissimilar* than would be expected by chance if they were identical. but are demonstrably not identical. In other words, dACC responses reflect partial integration across dimensions, not complete integration. We also observed putative signatures of comparison in these neurons, suggest that comparison does not require evaluation to be complete to proceed. These results suggest that dACC, even though it is normally placed late in the choice hierarchy, does not follow the completion of integration. Moreover, we believe these results are more consistent with distributed than with highly modular accounts of economic choice.

## RESULTS

### Behavior is informed by integrated value

Details of the *token risky choice task* are provided in the **Methods** and are illustrated in **Figure 1A** (see also Azab and Hayden, 2017 and 2018). On each trial, a macaque subject chose between two gambles. Gambles offered the possible gain or loss of virtual tokens. Tokens accumulated across trials; once subjects accrued six, they received a large water reward. Gambles consisted of two possible outcomes (which we denote “large” and “small”—the small one was most often negative or zero). Gambles appeared asynchronously; this staggered presentation allowed us to examine neuronal responses to the first gamble independent of the value of the second.

**Figure 1:**
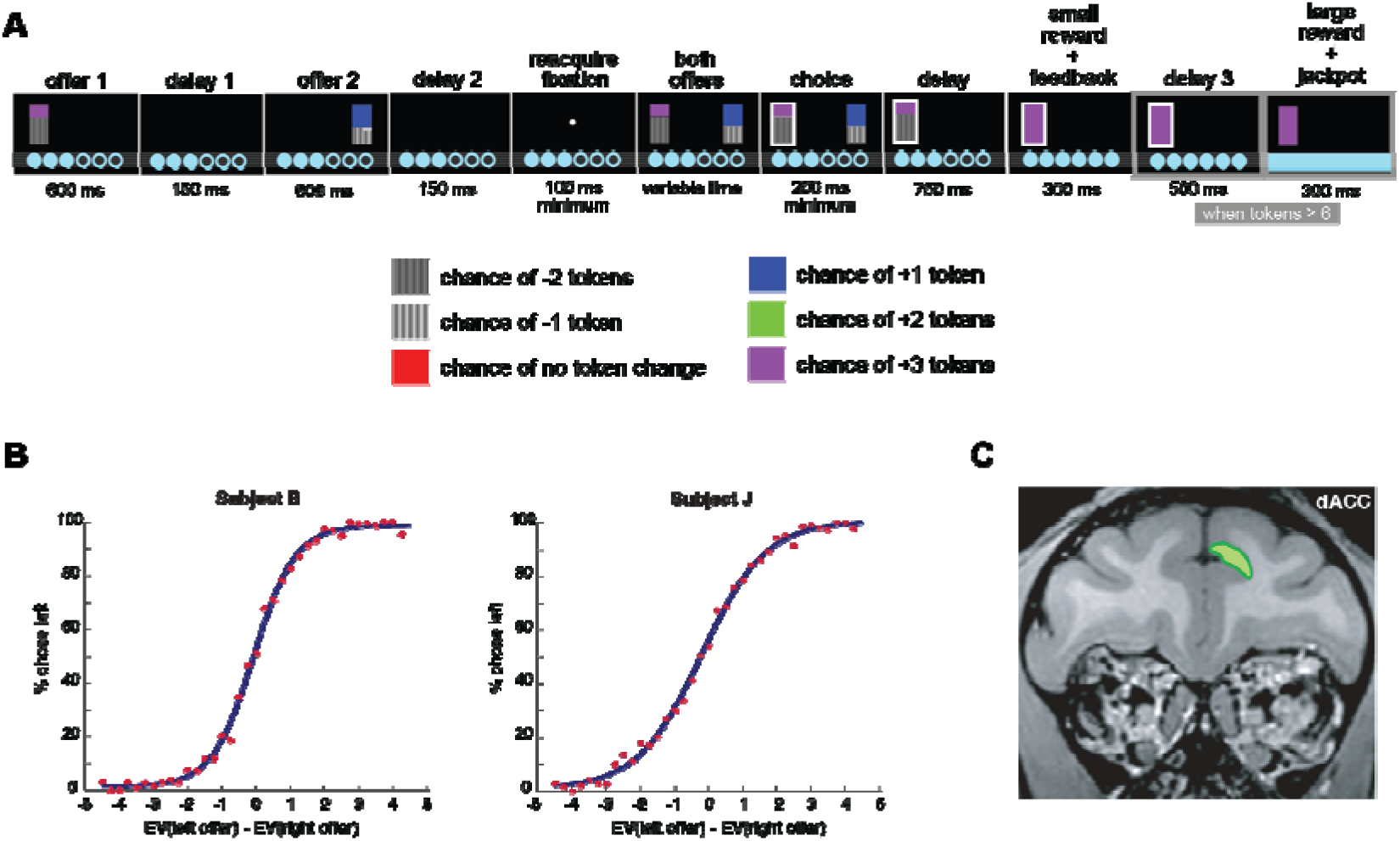
**A:** Example trial from token-gambling task. **B.** Behavior for each subject, fit to a sigmoid function. Subjects choose the left option more often as its value increases, as would be expected given understanding of the task. EV: expected value of gamble. **C:** Regions of interest (for exact coordinates, see Methods). Figure adapted with permission from Azab & Hayden, 2018.

#### Gamble values influence subjects’ choices

Details of the subjects’ behavior are provided elsewhere (Azab and Hayden, 2017; Azab & Hayden, 2018; Farashahi et al., 2018). Briefly, both subjects preferred the option with the higher expected value (subject B: 80.3%; subject J: 75.1%; two-sided binomial test: both P < 0.0001). Behavior was significantly influenced by all three components (the two outcomes and the probability) of both gambles (see **Table 1**).

**Table 1:** Subjects’ choices were influenced by all values characterizing both gambles. Average regression coefficients from a multiple logistic regression model of choice (left=1, right=0) against the variables in the regressor column, for each behavioral session. The distribution of regression coefficients across sessions for every behavioral variable listed deviated significantly from zero. A ‘loss’ within a gamble was always less than or equal to the win outcome of that gamble, and may or may not be a negative value. The magnitude of the regression coefficient indicated the extent to which this variable influenced choice, while its sign indicated whether higher values of this variable (positive vs. negative) favored choice of the left vs. right gamble, respectively. Adapted with permission from Azab & Hayden, 2018.

|  | Subject B (n = 66 sessions) |  | Subject J (n = 74 sessions) |  |
| --- | --- | --- | --- | --- |
| **Regressor** | Mean regression coefficient | Wilcoxon sign-rank Z-statistic (p-value) | Mean regression coefficient | Wilcoxon sign-rank Z-statistic (p-value) |
| Left offer – win | 1.40 | 7.06 (p = $1.64 \times 10^{-12}$) | 1.15 | 7.47 (p = $7.773 \times 10^{-14}$) |
| Left offer – loss | 0.486 | 6.97 (p = $3.26 \times 10^{-12}$) | 0.293 | 7.22 (p = $6.35 \times 10^{-13}$) |
| Left offer – probability of win | 5.12 | 7.06 (p = $1.64 \times 10^{-12}$) | 3.70 | 7.47 (p = $7.73 \times 10^{-14}$) |
| Right offer – win | -1.33 | -7.06 (p = $1.65 \times 10^{-12}$) | -1.33 | -7.47 (p = $1.67 \times 10^{-12}$) |
| Right offer – loss | -0.412 | -6.83 (p = 8.37 x 10<sup>-12</sup>) | -0.237 | -7.06 (p = 1.67 x 10<sup>-12</sup>) |
| Right offer – probability of win | -4.85 | -7.06 (p = 1.64 x 10<sup>-12</sup>) | -4.09 | -7.47 (p = 7.73 x 10<sup>-14</sup>) |

We first tested whether *choices* reflect the integrated values of the gambles or the components of each gamble independently. That is, whether the weighted product of outcome magnitudes and probabilities explained behavior over and above these gamble components individually. We fit choices to two multiple linear regression models, one including the three individual components of each gamble and another that included their interaction (that is, a multiplicative term). For both subjects, inclusion of the integrated value predictor improved model fit (Wilcoxon sign-rank test between Akaike Information Criterion, AIC, subject B: z=7.06; subject J: z=7.47; p<0.0001 for both subjects). These results provide important evidence that our subjects behaviorally integrate the values. In other words, they indicate that the final common pathway (the oculomotor control neurons) encode integrated value, and therefore indicate that the integration must occur before or at that point (Roesch and Olson, 2003; Roesch and Olson, 2004).

Burnham and Anderson recommend using a slightly different measure (AICc) when the ratio between the number of data points and model parameters is smaller than 40 (2004). Although this concern does not apply to our dataset (except in two sessions), the AICc criterion can safely be used in our datasets because it approaches AIC when that ratio gets larger. To that end, we repeated this analysis with AICc (rather than standard AIC). Our results remain qualitatively unchanged; a Wilcoxon sign-rank test suggests AICc is consistently higher for models not including integrated value predictors (Z-stat = 7.0087, p < 0.0001), indicating that models with integrated value predictors perform better. Moreover, the model including the integrated value regressors has a lower AICc measure for all behavior sessions.

### Responses of dACC neurons carry value-relevant information

We recorded 129 neurons from dACC in two subjects (subject B: 55 neurons, subject J: 74 neurons; see **Figure 1B** for region of interest). Basics of modulation in response to task-relevant variables are presented in detail in previous manuscripts and are not repeated here (Azab & Hayden, 2017 and 2018). We use the same epoch we used in our previous papers, a 500 ms window starting 100 ms after events. Example cells are shown in **Figure 2**. For the first example cell, lower low values result in higher firing rates (t-test on firing rates, p<0.001). For the second example cell, higher high values result in higher firing rates (p<0.005). For the third example cell, firing depended on the probability of the large outcome (p<0.001).

**Figure 2:**
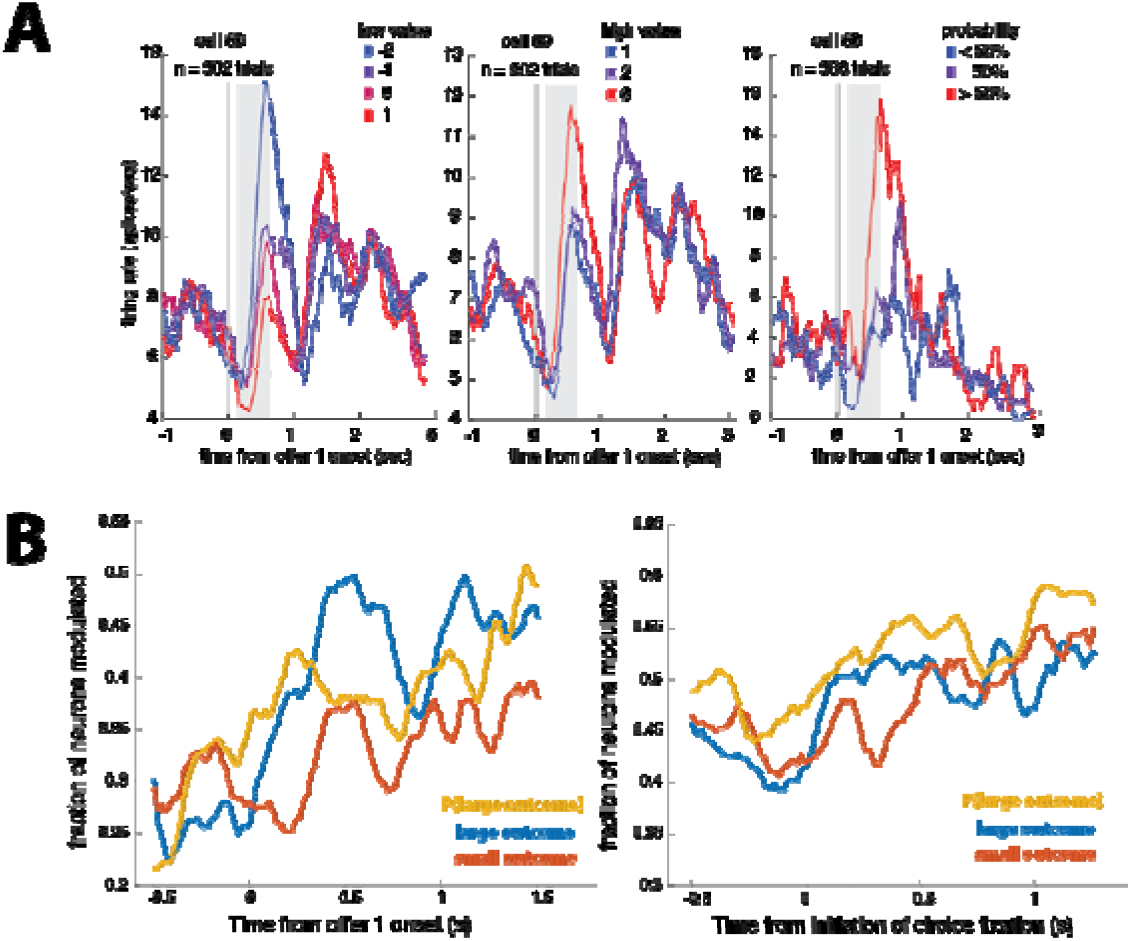
Neural responses to gamble components. **A:** Responses of example neurons to the small outcome (left), large outcome outcome (middle), and probability outcome (right) of the first gamble. Gray region: period of significant modulation. **B:** Proportion of neurons modulated by the components of the first gamble throughout the course of the trial. This fraction was computed by fitting a stepwise regression model to the normalized firing rate over a 500 ms sliding window against the components of the attended gamble (left) or the chosen gamble (right), along with other task-relevant variables (see Methods). Apparent effects before time zero reflect the width of the analysis window.

Responses of 17.05% of neurons were affected by the size of the large outcome (n=22/129, one-sided binomial test: p<0.0001). Responses of 20.2% were affected by the probability of that outcome (n=26/129, p<0.0001), and 8.53% were affected by the small outcome (n=11/129, p=0.059) of the same gamble in the first epoch. Note that the lack of a significant effect for the small outcome raises the possibility that dACC simply does not encode this parameter. However, more sensitive tests do show an effect. Specifically, we performed a stepwise linear regression model; a significant fraction of neurons showed dependence on the small outcome in both epochs (epoch 1: 34.1%, n = 44/129, P < 0.0001; epoch 2: 55.0%, n = 71/129, two-sided binomial test: P < 0.0001).

In our sample, the firing rate of 27.1% (n=35/129) of neurons encoded the expected value of the first offer during the epoch it appeared (this ratio is significant; 2-sided binomial test, p < 0.0001). During the second epoch, 23.3% (30/129) of neurons encoded the offer 1 value (p < 0.0001). This finding suggests that the offer 1 value was retained, presumably in working memory, even when it was no longer on the screen. During the second epoch, 16.3% (21/129) of neurons encoded the value of the second offer (p < 0.0001).

### dACC neurons use partially integrated formats to encode gamble outcomes

We have used the term *ensemble coding format* (or just *format*) to refer to the vector of tuning coefficients for all recorded neurons (Strait et al., 2014; Wang and Hayden, 2017). Specifically, we used betas from the full regression as our tuning coefficients. We then correlate formats across conditions to determine how they relate. Bayesian regression allows us to draw samples from the likely format (rather than the single estimate conventional regression provides, Gelman et al., 2013). All correlations below use Spearman (i.e. rank-order) correlation, which is more robust to outliers than Pearson correlation, and is thus a more conservative method. This analysis approach has a good deal of conceptual resemblance to representational similarity analysis (RSA; Kriegeskorte et al., 2008); the main practical difference is that it uses parameter tuning (specifically, regression weights) rather than raw response (e.g. firing rates), making it more appropriate for the questions we want to ask here (see **Discussion**).

It is important to note how correlation of betas (the focus of this study) differs from comparison of size of betas (which this study does not deal with.) We could imagine for example that behavior depends on an unweighted (equal) sum of dimensions, while firing rate depends 10x on probability relative to stakes. To perform that analysis, we would compare the magnitudes of the regression coefficients associated with large and small outcomes—comparable coefficient magnitudes would suggest equal weighting of both outcomes. However, that is not the question we are concerned with in this manuscript. Mathematically, that question is about the relative strength of tuning, or the absolute slope of the line relating the betas for stakes and probability relative to behavior—that is, the *correlation* between these slopes rather than a comparison of their magnitude. A difference in the betas (such as could be found with a t-test) would indicate that the outcomes are not equally weighted. Instead what we show is that the variables are positively but not perfectly correlated (using a correlation).

We found that ensemble coding formats for large and small outcomes of each offer are positively correlated (r=0.28, p<0.001, 1000 samples). That is, our population uses overlapping (i.e. more similar than expected by chance) neural codes to signal the values of the large and small outcomes; this is a signature of cross-dimensional integration (Strait et al., 2014; Blanchard et al., 2015).

But does it reflect a *full* cross-dimensional integration? Mathematically speaking, why is the measured correlation coefficient less than 1.0? It could reflect intermediate levels of integration or it could reflect complete integration that appears intermediate due only to measurement noise. (Recall that neurons are inherently noisy, so we can be certain that such noise is a major factor). To differentiate these possibilities, we repeated the same analysis on a shuffled dataset (where the values of the two variables of interest are shuffled *within* trial, see **Methods**). This tells us the maximum correlation observable between the encoding of two variables given the noise in our data, which we call the *ceiling correlation*.

The ceiling correlation for large and small outcomes in the first epoch was r=0.45 (standard deviation = 0.047; **Figure 3A**). This result indicates that noise is high in our dataset (as it is in most neural recordings) and that good deal of the explanation for the low correlation we observed is simply noise. However, our observed correlation (r = 0.28, see above) is nonetheless significantly lower than this value (p=0.001, bootstrap significance test, 1,000 permutations). Thus, although large and small outcomes are encoded in similar formats, these formats are not identical, as one would expect if dACC outputs reflected a completed value computation. Mathematically speaking, the difference in formats necessarily reflects a qualitative (non-affine) difference in tuning—a consistent gain shift or baseline shift would not be detected by our method and would not generate a correlation significantly different from ceiling. In other words, dACC neurons show evidence simultaneously of vectors corresponding integration and non-integration, and these vectors are combined in our sample of neurons.

**Figure 3:**
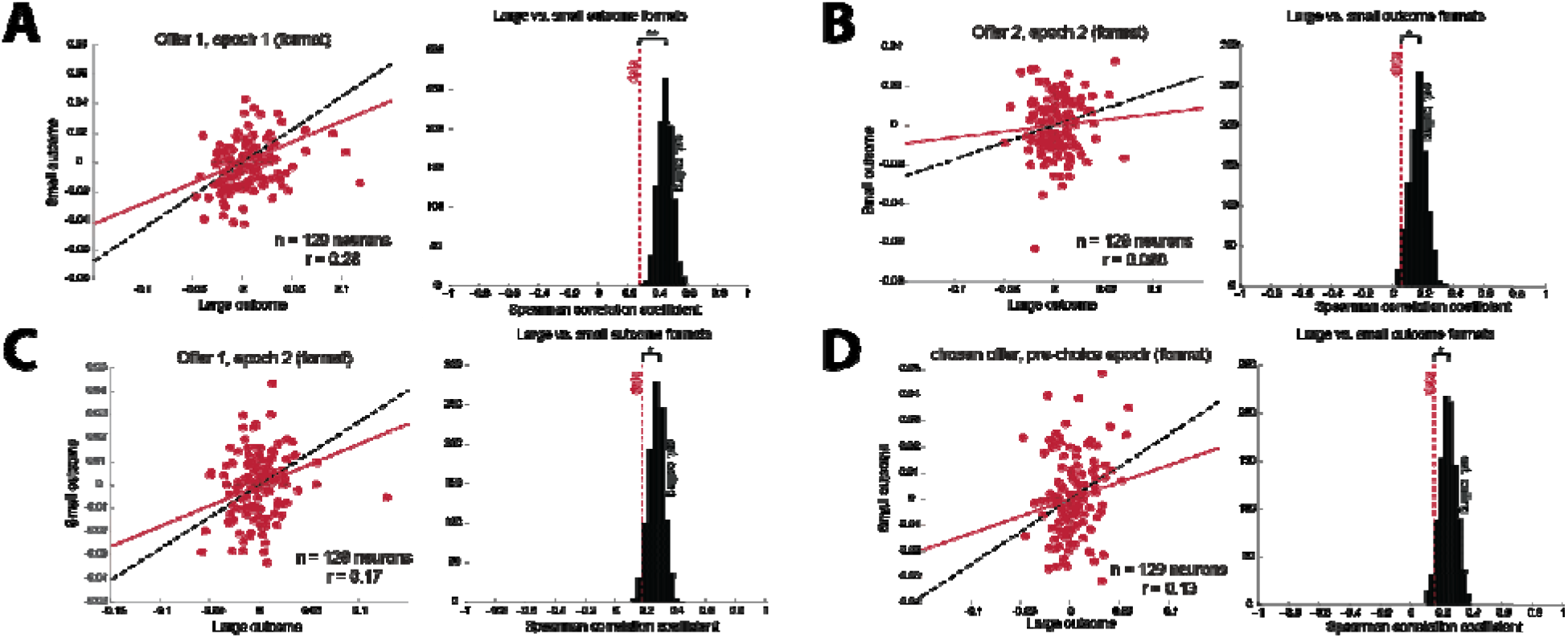
Large and small outcomes are encoded in partially-separable formats in the recorded population. **A:** Correlation between formats of large and small outcomes for the first gamble in the first epoch. (left) Scatter-plot showing the correlation between regression coefficients for large and small outcomes for each neuron. Red line indicates the positive correlation between regression coefficients. Shaded red region indicates the 95% credible interval on the data correlation. Black dashed line indicates the estimated ceiling correlation between these variables in this epoch. (right) Results of permutation test. Black histogoram indicates the distribution of the estimated ceiling correlation from permuted data. Red dashed line indicates degree of correlation observed in the data. **B:** Correlation between variable formats for the second gamble in the second epoch. **C.** Correlation between variable formats for the first gamble in the second epoch. **D:** Correlation between variable formats for the chosen gamble in the pre-choice epoch. *: p < 0.05, **p < 0.01.

We found a similar pattern when considering the second gamble in the *second offer epoch* (t=100 to 600 ms after the second offer appears). Specifically, although we did not find evidence of integration across the two dimensions (r=0.059, p=0.060, compared to zero), we found that whatever integration existed was less than the ceiling correlation value (r=0.17; p=0.028; Figure 3B). Thus, it appears that dACC does not fully integrate either offer. (Note that, because it is a null result, the lack of integration observed for the second offer does not prove no integration occurred, although it is consistent with that possibility.)

We next asked whether full integration occurs when the information about the gamble is transferred to working memory. It appears not. The coding of the first offer during the second offer epoch (when it was no longer on screen) resulted in the same pattern: partial but not complete integration (r=0.18, p<0.001 for comparison with zero. Ceiling correlation was r=0.27, which was larger than the observed value, p=0.031; **Figure 3C**).

### dACC continues to show partial integration around the time of choice

We next examined the 500 ms immediately preceding choice fixation (*pre-choice epoch*). Whereas above we focused on offered gambles, here we focused on the *chosen gamble*, with the reasoning that identifying an option for choice would potentially result in full integration. The formats for integration of the two stakes is, indeed, positive (r=0.16, p<0.001). However, as like in previous epochs, this correlation is less than ceiling (r=0.25, p=0.047; Figure 3D). These results thus show that separation in large- and small-outcome formats persists at least until right before choice is indicated.

### Formats used to encode stakes and probabilities are consistent with full integration for those dimensions

For the first gamble in the first epoch, we find that the large outcome and its probability were encoded in similar formats (r=0.54, p<0.001). This degree of correlation was not significantly less than our estimated ceiling (ceiling r=0.58, p=0.135; Figure 4A). While we cannot definitively conclude that there is full integration of these variables, the difference between these results and the previous ones is striking. Indeed, for the first offer in the second epoch (i.e. when it is in working memory), large outcomes and probabilities are also encoded in similar formats (r=0.44, p<0.001) that are no different from ceiling (r=0.46, p=0.39; Figure 4B). For the second offer, large outcomes and their probabilities are also encoded in similar formats (r=0.37, p<0.001), and again no different from ceiling (r=0.41, p=0.22; Figure 4C). The chosen gambles’ large outcomes and probabilities are encoded in similar formats in the 500 ms preceding choice (r=0.49, p<0.001), and again no different from ceiling (r=0.47, p=0.63; Figure 4D). Although the lack of a difference is not sufficient to prove identity, the matching patterns for all four conditions, and their striking differences with the stakes analyses above, suggest that large stakes and probability may be fully integrated in dACC.

**Figure 4:**
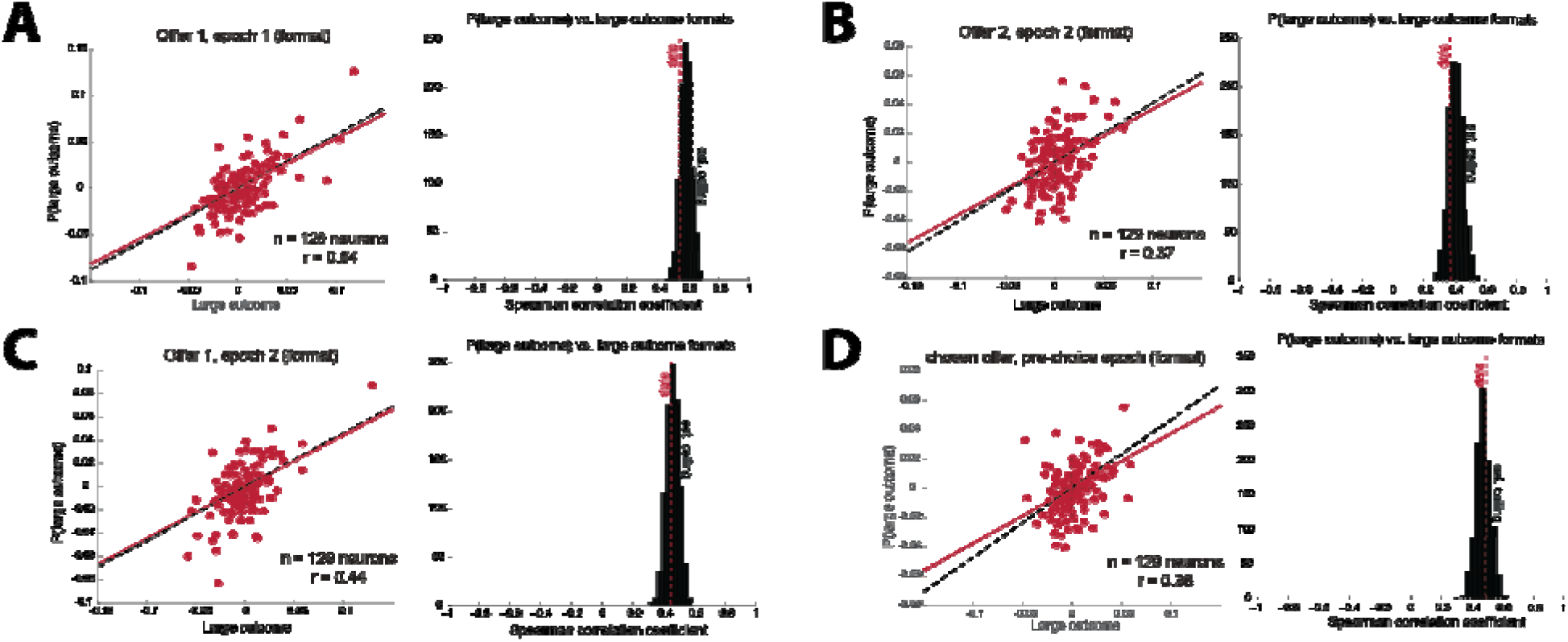
Large outcomes and their probabilities are encoded in integrated formats in the recorded population. **A:** Correlation between formats of large outcomes and probabilities for the first gamble in the first epoch. (left) Scatter-plot showing the correlation between regression coefficients for large outcomes and probabilities for each neuron. Red line indicates the positive correlation between regression coefficients. Shaded red region indicates the 95% credible interval on the data correlation. Black dashed line indicates the estimated ceiling correlation between these variables in this epoch. (right) Results of permutation test. Black histogoram indicates the distribution of the estimated ceiling correlation from permuted data. Red dashed line indicates degree of correlation observed in the data. **B:** Correlation between variable formats for the second gamble in the second epoch. **C.** Correlation between variable formats for the first gamble in the second epoch. **D:** Correlation between variable formats for the chosen gamble in the pre-choice epoch.

These results are important because they indicate that the lack of integration observed for large and small stakes is not a consequence of our task design, of the psychology of our subjects, or of a limitation in our analysis approach. It appears that dACC is capable of fully integrating some variables and maintaining some separation in others, and our methods are capable of detecting this partial integration when it occurs.

This final point is further emphasized by our next analysis. In the first epoch, the small outcomes and probabilities were encoded in similar formats (r=0.11, p=0.002), which are less correlated than ceiling (r=0.30, p=0.001). Results are not conclusive for the same gamble in the second epoch: there is no correlation between the formats for small outcomes and large-outcome probabilities (r=0.050, p=0.11), but this correlation is no different from our estimated ceiling, which is very low (r=0.13, p=0.057). The small outcome and large-outcome probability of the second gamble are encoded in weakly-correlated formats in the second epoch (r=0.063, p=0.049), and this correlation is no less than our estimated ceiling (also low, ceiling r=0.13, p=0.12). The low stakes and probability components of the chosen gamble were encoded in positively-correlated formats in the pre-choice epoch (r=0.17, p<0.001); no less than ceiling (r=0.15, p=0.65). The lack of full integration between small outcomes and probabilities in the first epoch (which is highly significant and survives Bonferroni correction) highlights the qualitatively different way in which large and small offer outcomes are treated.

## DISCUSSION

We examined the neural encoding of the values of risky offers in dACC. We made use of a recently-developed statistical method that can disambiguate variable encodings that are partially correlated from ones that are fully correlated but different due solely to noise (Azab and Hayden, 2017). We find, first, that while dACC neurons encode both large and small prospective outcomes of individual gambles on the screen and in working memory, the population tuning formats for the two outcomes are only partially correlated—meaning neither orthogonal nor collinear.

The “mid-evaluation” signal we observe is a type that is predicted by hierarchical (sometimes called serial) models of value, which involve multiple small steps leading from a space of options to one of actions rather than a single value estimation locus (Yoo & Hayden, 2018). In such models, reward and value processing occur along an anatomical hierarchy and computation within that hierarchy implements a gradual transformation from stimulus information to motor commands, rather than a series of categorical steps (Hunt et al., 2012; Hunt & Hayden, 2017; Cisek, 2012; Chen & Stuphorn, 2015; Cisek & Kalaska, 2010).

More specifically, we have proposed that the relationship between different hierarchical levels may be one of untangling—of rotating representations into a format more usable for effector systems, while retaining information (Yoo and Hayden, 2018; DiCarlo et al.. 2012). The untangling theory, which is based on how the visual stream works, proposes that each anatomical stage untangles information from the earlier stage and allows for intermediate amounts of untangling. In the domain of economic choice, the tangled format involves the two non-integrated dimensions and the untangled format is an integrated value representation. Our study provides tentative support for this idea.

The first major effect we observe suggests that dACC follows the initiation of value integration but precedes its completion. Together these results suggest that dACC cannot be assigned either a purely pre- or post-evaluation stage in choice. Moreover, in conjunction with other results showing evidence for value comparison in dACC (e.g. Azab & Hayden, 2017; Azab & Hayden, 2018; Klein-Flugge & Bestmann, 2012; Hare et al. 2011; Wunderlich et al., 2009; Rangel & Hare, 2010), these findings indicate that value integration does not need to be complete before value comparison begins (Hunt et al., 2014).

These results complement our recent findings using this dataset (Azab & Hayden, 2017). In that study, we concluded that dACC occupies a middle role in the *comparison* of values. Our complementary results here suggest that dACC occupies a middle role in the *computation* of values. Juxtaposed, the two results have a larger implication: that the completion of integration is not essential to the initiation of comparison. That is, it may be impossible, even in theory, to draw a line, either in anatomical space or in time, between evaluation and comparison stages of choice (Yoo & Hayden, 2018).

The identification of abstract amodal value signals is a desideratum of neuroeconomics (Shizgal, 1997; Padoa-Schioppa, 2011; O’Doherty, 2014; Montague & Berns, 2002; Levy & Glimcher, 2012). Our results suggest that, if these exist, they are most likely to occur only downstream of dACC. We envision two major possibilities about this site of value representation. First, value representations may only be complete in the premotor or motor system (see Cisek, & Kalaska 2010, Cisek, 2007, and Thura & Cisek, 2016 for related arguments). Second, that may be no single location at which value is represented abstractly and completely (Chen & Stuphorn, 2015; Cisek, 2012; Yoo & Hayden, 2018; Hunt et al., 2015; Balasubramani et al., 2018).

Ours is not the first study to consider representations of multi-attribute offers (O’Neill & Schultz, 2018; Raghuraman & Padoa-Schioppa, 2014; Kennerley et al., 2009; Strait et al., 2014; Blanchard et al., 2015; Padoa-Schioppa & Assad, 2006; Hunt et al., 2015). A broad finding from these studies is that neurons in several regions show some overlap in the way they encode the multiple dimensional determinants of value. The major difference is that we show that integration of attributes into value representations is not an all-or-none phenomenon. The distinction between attributes that are not integrated vs. somewhat integrated may not be meaningful for readout, since this information is extractable in either case—future work is needed to know what information is read out of dACC responses.

Our data touch on but do not provide dispositive evidence relating to a larger discussion in the literature – whether choice occurs without full integration (Busemeyer et al., 2019). In our view, the most likely answer is “sometimes yes and sometimes no”. Indeed, we have other work suggesting that in some other cases, full integration is not supported by behavior. However, in the specific case of this task, there is strong behavioral evidence of integration.

We have proposed that what we call a value may be better thought of as evidence in favor of the proposition that the option will be chosen (Azab & Hayden, 2017; see also Hayden & Moreno-Bote, 2018). From this perspective, there is no reason to expect that valuation must be complete before comparison can begin. Each dimension can provide its own evidence in favor or against the choice of an option. That evidence can be summed in one specific place or it can be fed into a slowly evolving action plan without ever being integrated in one place. We and others have previously argued that the analogy to bee swarm decision-making is helpful for understanding choice (Hunt & Hayden, 2017; Eisenreich et al., 2017; Couzin, 2009; Passino et al., 2008; Seeley et al., 2012; Pirrone et al., 2018). Individual bees evaluate hive sites in an extremely noisy fashion (Seeley, 2009; Seeley & Buhrman, 1999). Indeed, an individual scout may only assess one or two of the several dimensions along which hives vary. Even if no individual bee performs a full cross-dimensional integration, the hive as a whole can still make a good decision—without ever computing anything like value—through a quorum sensing procedure. And the hive as a whole takes several days to select a hive site. In the middle of this time, the hive may be said to have made a partially-complete evaluation process.

Similarly, some *cognitive* (rather than neural) models of value-based decision-making do not make this assumption of discrete, serial stages; ordinal within-attribute comparisons oftentimes contribute to the construction of each option’s value. In this framework, comparison would, in some sense, *precede* evaluation, and all value signals are entirely relative (Noguchi & Stewart, 2018; Busemeyer et al., 2019). These models—also ones of value-based decision-making, albeit at a different level of granularity—clearly demonstrate the importance of integrating different levels of analysis when exploring the inner workings of a cognitive process, and the many ways that a system (biological or otherwise) could potentially instantiate it.

## ACKNOWLEDGEMENTS

We would like to thank Meghan Castagno, Marc Mancarella, and Caleb Strait for assistance with data collection. This research was funded by a National Science Foundation grant (grant number: BCS1253576) received by BYH, and a National Institutes of Health grant (grant number: DA038615) received by BYH

## MATERIALS AND METHODS

The data used here were analyzed and summarized in other manuscripts (Strait et al., 2015; Azab & Hayden, 2017; Azab & Hayden, 2018; Farashahi et al., 2018).

### Surgical Procedures

All procedures were approved by the University Committee on Animal Resources at the University of Rochester and were designed and conducted in compliance with the Public Health Service’s Guide for the Care and Use of Animals. Two male rhesus macaques (*Macaca mulatta:* subject B age 5y. 7mo.; subject J age 6y. 7mo. at the start of recording) served as subjects in both studies. We used standard procedures, as described previously (Strait et al., 2014). A small prosthesis for holding the head was used. Animals were habituated to laboratory conditions and then trained to perform oculomotor tasks for liquid reward. A Cilux recording chamber (Crist Instruments, Hagerstown, Maryland, USA) was placed over the dACC and attached to the calvarium with ceramic screws. Appropriate anaesthesia was used at all times; induction was performed with ketamine and isoflurane was used for maintenance. For surgical induction, we used 10–15 mg/kg of ketamine, 0.25 mg/kg of midazolam, and 2–4 mg/kg of propofol. For maintenance, we used isoflurane, ad lib level, set depending on active monitoring procedure. For systemic antibiotics, we used cefazolin and for topical application, we used standard veterinary triple antibiotic. For analgesics, we used meloxicam, and, when judged necessary by veterinary staff, buprenorphine. Post-operative care included close monitoring and restoration of fluid intake. Animals received appropriate analgesics and antibiotics after all procedures. Position was verified by magnetic resonance imaging with the aid of a Brainsight system (Rogue Research Inc., Montreal, Quebec, Canada). Throughout both behavioral and physiological recording sessions, the chamber was kept sterile with regular antibiotic washes and sealed with sterile caps.

### Recording Site

We approached dACC through a standard recording grid (Crist Instruments). We defined dACC according to the Paxinos atlas (Paxinos et al, 2000). Roughly, we recorded from a ROI lying within the coronal planes situated between 29.50 and 34.50 mm rostral to interaural plane, the horizontal planes situated between 4.12 to 7.52 mm from the brain’s dorsal surface, and the sagittal planes between 0 and 5.24 mm from medial wall (Figure 1B). Our recordings were made from a central region within this zone. We confirmed recording location before each recording session using our Brainsight system with structural magnetic resonance images taken before the experiment. Neuroimaging was performed at the Rochester Center for Brain Imaging, on a Siemens 3T MAGNETOM Trio Tim using 0.5 mm voxels. We confirmed recording locations by listening for characteristic sounds of white and gray matter during recording, which in all cases matched the loci indicated by the Brainsight system.

### Electrophysiological Techniques

Single electrodes (Frederick Haer & Co., Bowdoin, Maine, USA; impedance range 0.8 to 4 MU) were lowered using a microdrive (NAN Instruments, Nazaret Illit, Israel) until waveforms of between one and three neuron(s) were isolated. Individual action potentials were isolated on a Plexon system (Plexon, Inc., Dallas, Texas, USA). Neurons were selected for study solely on the basis of the quality of isolation; we never pre-selected based on task-related response properties. All collected neurons for which we managed to obtain at least 250 trials were analyzed.

### Eye Tracking and Reward Delivery

Eye position was sampled at 1,000 Hz by an infrared eye-monitoring camera system (SR Research, Ottawa, Ontario, Canada). Stimuli were controlled by a computer running Matlab (Mathworks) with Psychtoolbox (Brainard, 1997) and Eyelink Toolbox (Cornelissen, Peters & Palmer, 2002). Visual stimuli were colored rectangles on a computer monitor placed 57 cm from the animal and centered on its eyes. A standard solenoid valve controlled the duration of juice delivery. The relationship between solenoid open time and juice volume was established and confirmed before, during, and after recording.

### Behavioral Task

Subjects performed a two-option gambling task. The task was conceptually similar to risky choice tasks we have used previously (Strait et al., 2014), with two major differences: (1) monkeys gambled for virtual tokens—rather than liquid—rewards, and thus (2) outcomes could be losses as well as wins. Previous training history for these subjects included a set shifting task (Sleezer et al., 2016), two simple choice tasks (Blanchard et al., 2014; Heilbronner and Hayden, 2016; Yoo and Hayden, 2019), an attentional task (Hayden and Gallant, 2013), and a pursuit task (Yoo et al., 2020).

Two offers were presented on each trial. Each offer was represented by a rectangle 300 pixels tall and 80 pixels wide (11.35° of visual angle tall and 4.08° of visual angle wide). 20% of options were safe (100% probability of either 0 or 1 token), while the remaining 80% were gambles. Safe offers were entirely red (0 tokens) or blue (1 token). The size of each portion indicated the probability of the respective reward. Each gamble rectangle was divided horizontally into a top and bottom portion, each colored according to the token reward offered. Gamble offers were thus defined by three parameters: two possible token outcomes, and probability of the top outcome (the probability of the bottom was strictly determined by the probability of the top). The top outcome was 10%, 30%, 50%, 70% or 90% likely on gamble offers.

Six initially unfilled circles arranged horizontally at the bottom of the screen indicated the number of tokens to be collected before the subject obtained a liquid reward. These circles were filled appropriately at the end of each trial, according to the outcome of that trial. When 6 or more tokens were collected, the tokens were covered with a solid rectangle while a liquid reward was delivered. Tokens beyond 6 did not carry over, nor could number of tokens fall below zero.

On each trial, one offer appeared on the left side of the screen and the other appeared on the right. Offers were separated from the fixation point by 550 pixels (27.53° of visual angle). The side of the first offer (left and right) was randomized by trial. Each offer appeared for 600 ms and was followed by a 150 ms blank period. Monkeys were free to fixate upon the offers when they appeared (and in our observations almost always did so). After the offers were presented separately, a central fixation spot appeared and the monkey fixated on it for 100 ms. Following this, both offers appeared simultaneously and the animal indicated its choice by shifting gaze to its preferred offer and maintaining fixation on it for 200 ms. Failure to maintain gaze for 200 ms did not lead to the end of the trial, but instead returned the monkey to a choice state; thus, monkeys were free to change their mind if they did so within 200 ms (although in our observations, they seldom did so). A successful 200 ms fixation was followed by a 750 ms delay, after which the gamble was resolved and a small reward (100 μL) was delivered—regardless of the outcome of the gamble—to sustain motivation. This small reward was delivered within a 300 ms window. If 6 tokens were collected, a further delay of 500 ms was followed by a large liquid reward (300 μL) within a 300 ms window, followed by a random inter-trial interval (ITI) between 0.5 and 1.5 s. If 6 tokens were not collected, subjects proceeded immediately to the ITI.

Each gamble included at least one positive or zero-outcome, ensuring that every gamble carried the possibility of a win (or at least no change in tokens). This decreased the number of trivial choices presented to subjects, and maintained motivation.

### Statistical Methods

**Peri-stimulus time histograms (PSTHs)** were constructed by aligning spike rasters to the presentation of the first offer and averaging firing rates across multiple trials. Firing rates were calculated in 20 ms bins but were generally analyzed in longer epochs. For latency analyses, we used PSTHs with firing rates calculated in 5 ms bins to grant us finer resolution to detect shorter latencies.

Firing rates were **normalized** by subtracting the mean and dividing by the standard deviation of the neuron’s entire PSTH (i.e. z-scoring).

We tested for correlations between variables of interest, since these could produce spurious results in our format and population correlation analyses. Although gamble values were chosen independently, safe gambles (where the large and small outcome were of the same value) introduced a spurious correlation between large and small outcomes. We thus exclude trials where the gamble of interest was a safe gamble, as we have done in previous manuscripts (refs).

We tested for single-unit modulation using a **multiple generalized linear regression model**, including the following task-relevant variables: large outcomes, small outcomes, and probability of the large outcomes for gambles 1 (and two, if that gamble had been presented), the number of tokens collected as of the beginning of the trial, and the side the first offer appeared on. We also included the side of the chosen offer for analyses in the pre-choice and post-choice epochs. We use the same variables to fit stepwise linear regression models to assess small-outcome modulation. We used stepwise linear regression as a more sensitive method to determine whether small outcomes were, in fact, encoded. To do this, we used the stepwiselm function in Matlab, with a base model of linear terms and default settings. The predictors used for this procedure were the small outcome, the large outcome, and the probability of the large outcome. These values were included for only the first offer only until offer 2 appeared, and for both offers after that.

**Analysis epochs** were chosen *a priori*, before data analysis began, to align with epochs we have chosen for previous studies (Strait et al., 2014; Azab & Hayden, 2017; Azab & Hayden, 2018). The first and second offer epochs were defined as the 500 ms epoch beginning 100 ms after the offer was presented. The pre-choice epoch was the 500 ms epoch before choice was indicated using an express saccade. The post-choice epoch was the 500 ms epoch immediately after choice fixation was complete, and before reward feedback was revealed. These epochs were chosen before data collection began; indeed, they originally were selected for a previous study on a different brain region, and to account for hypothesized delays in neural responding (Strait et al., 2014). We maintained the same epochs here as a mechanism to reduce the likelihood of p-hacking.

All **fractions of neurons** were tested for significance using a one-sided binomial test. When testing for significant encoding in the population, we use an alpha level of 0.05 to indicate chance (i.e. the number of neurons that would exhibit significant modulation at random).

### Format and population correlation analyses

Analyses were performed in the following manner. We use **beta correlation analyses** to assess whether neurons represented two variables (or the same variable at different time periods) using similar / orthogonal / opposing formats, in overlapping / orthogonal / distinct populations. To do this, we first found the regression coefficient associated with the variable of interest per neuron. We estimate this using a multiple linear regression model including the same variables detailed above (for spatial location analyses, we split trials by the side of offer appearance, and thus do not include that variable in our regression model). We then combined the regression coefficients associated with the variable of interest into a vector of the same length as the number of neurons in our sample. This vector indicates the relative strength (after normalizing) and direction of modulation for each individual neuron in the population, in response to a particular variable in a particular epoch. We call this the population “format”. We compared different formats by finding the Spearman correlation coefficient between them.

Correlations of signed and unsigned regression coefficients can provide very different types of information about the population. Correlations of signed coefficients provide information about the relationship between *coding formats* in a population, and are good for identifying whether neurons encode in the same format in different conditions; correlations of unsigned coefficients provide information about the relationship between *strengths*, and are good for identifying whether similar or distinct populations of cells are used.

Note that this approach provides a more sensitive method of examining population properties than conventional approaches, which involve determining which cells cross a significance threshold and then using those as focal cells for further analyses. By taking on all cells regardless of their response, our method accomplishes two things. First, it uses all available information, even that information that is not sufficient by itself to achieve significance. Second, it avoids introducing a hard categorical boundary, which can introduce false positives in data (Blanchard et al., 2017; Maxwell & Delaney, 1993).

We extend this method to account for the noise inherent in estimating each neuron’s encoding of each variable of interest, which existing methods do not account for (Azab & Hayden, 2017). We used a Bayesian regression to obtain a probabilistic distribution over each regression coefficient for each neuron (rather than an individual coefficient estimate per neuron; this is akin to taking into account the confidence interval on each regression coefficient estimate). We sampled 1,000 regression coefficients from this distribution for each neuron, to obtain a probability distribution of (1,000) potential formats for the population, for each of two task conditions. We then performed the correlation analyses on each of these samples across task conditions, thus generating a probability distribution of (1,000) correlation coefficients. This is a more robust estimate of the correlation between formats, as it takes into account the uncertainty inherent in estimating any individual regression coefficient, and allows us to view the spread of the distribution of this correlation when this significant source of noise is taken into account.

The Spearman correlation coefficient between signed regression coefficients indicated whether variables were represented in a similar *format* i.e. directionality of tuning across the population. (Note that we used Spearman instead of Pearson because it is more robust to outliers and is therefore more statistically conservative). A positive correlation indicated a preservation of directionality, while a negative correlation suggested variables were represented in opposing directionality of firing rate modulation. No correlation suggests orthogonal formats, but we draw no strong conclusions from these.

Similarly, the correlation coefficient between unsigned regression coefficients indicated whether similar neuronal populations tended to be involved in encoding the two variables in question, regardless of their direction of modulation. A positive correlation indicated overlapping populations, while a negative correlation indicated separate ones. A lack of correlation suggests orthogonal populations (i.e. encoding one variable does not affect the neuron’s likelihood of encoding the other variable), although given the lack of interpretability of null results, it does not constitute evidence for lack of true correlation.

It is important to note that the analysis method we use here is rooted in previous analysis techniques used by our lab and others. The closest relative is Representational Similarity Analysis (RSA; Kriegeskorte, Mur & Bandettini, 2008; Hunt et al., 2018). In RSA, neural representations (vectors of neural responses) associated with different behavioral conditions are compared and their similarity characterized to facilitate comparison with different data sets, sometimes collected using different means. Rather than comparing raw neural responses, though, our method relies on comparing regression coefficients; this allows us to hone in on the signal associated with a particular variable, allowing us to compare the neural responses to different variables at the same point in time.

### Shuffling approach to estimate ceiling correlation

Our approach was straightforward. Labels were shuffled and then we repeated the analysis to obtain new beta coefficients. We repeated this analysis within cells rather than across cells and within trials rather than across trials. We did this by randomly reassigning labels for large and small outcome within each trial. This approach maintains the relationship between firing rate and outcome on each trial, and only randomizes the variable of interest. If as a result there is no difference in that encoding, the coefficients shouldn’t change and the correlation should look the same as the data; it would suggest that the neuron is representing those two values, regardless of which is which.

## REFERENCES

Azab, H., & Hayden, B. Y. (2017). Correlates of decisional dynamics in the dorsal anterior cingulate cortex. PLoS biology, 15(11), e2003091.

Azab, H., & Hayden, B. Y. (2018). Correlates of economic decisions in the dorsal and subgenual anterior cingulate cortices. European Journal of Neuroscience, 47(8), 979–993.

Balasubramani, P.P., Moreno-Bote, R., and Hayden, B.Y. (2018). Using a simple neural network to delineate some principles of distributed economic choice. Front. Comput. Neurosci. 12, 22.

Blanchard, T. C., Hayden, B. Y., & Bromberg-Martin, E. S. (2015). Orbitofrontal cortex uses distinct codes for different choice attributes in decisions motivated by curiosity. Neuron, 85(3), 602–614.

Blanchard, T. C., Piantadosi, S., & Hayden, B. Y. (2017). Robust mixture modeling reveals category-free selectivity in reward region neuronal ensembles. Journal of neurophysiology.

Blanchard, T. C., & Hayden, B. Y. (2014). Neurons in dorsal anterior cingulate cortex signal postdecisional variables in a foraging task. Journal of Neuroscience, 34(2), 646–655.

Blanchard, T. C., Wolfe, L. S., Vlaev, I., Winston, J. S., & Hayden, B. Y. (2014). Biases in preferences for sequences of outcomes in monkeys. Cognition, 130(3), 289–299.

Bowman, N. E., Kording, K. P., & Gottfried, J. A. (2012). Temporal integration of olfactory perceptual evidence in human orbitofrontal cortex. Neuron, 75(5), 916–927.

Brainard, D. H., & Vision, S. (1997). The psychophysics toolbox. Spatial vision, 10, 433–436.

Burnham, K. P., & Anderson, D. R. (2004). Multimodel inference: understanding AIC and BIC in model selection. Sociological methods & research, 33(2), 261–304.

Busemeyer, J. R., & Townsend, J. T. (1993). Decision field theory: a dynamic-cognitive approach to decision making in an uncertain environment. Psychological review, 100(3), 432.

Busemeyer, J. R., Gluth, S., Rieskamp, J., and Turner, B. M. (2019). Cognitive and Neural Bases of Multi-Attribute, Multi-Alternative, Value-based Decisions. Trends in Cognitive Sciences 1873

Chau, B. K., Kolling, N., Hunt, L. T., Walton, M. E., & Rushworth, M. F. (2014). A neural mechanism underlying failure of optimal choice with multiple alternatives. Nature neuroscience, 17(3), 463.

Chen, X., & Stuphorn, V. (2015). Sequential selection of economic good and action in medial frontal cortex of macaques during value-based decisions. Elife, 4, e09418.

Cisek, P. (2007). Cortical mechanisms of action selection: the affordance competition hypothesis. Philosophical Transactions of the Royal Society of London B: Biological Sciences, 362(1485), 1585–1599.

Cisek, P. (2012). Making decisions through a distributed consensus. Current opinion in neurobiology, 22(6), 927–936.

Cisek, P., & Kalaska, J. F. (2010). Neural mechanisms for interacting with a world full of action choices. Annual review of neuroscience, 33, 269–298.

Cornelissen, F. W., Peters, E. M., & Palmer, J. (2002). The Eyelink Toolbox: eye tracking with MATLAB and the Psychophysics Toolbox. Behavior Research Methods, Instruments, & Computers, 34(4), 613–617.

Couzin, I. D. (2009). Collective cognition in animal groups. Trends in cognitive sciences, 13(1), 36–43.

David, S. V., & Hayden, B. Y. (2012). Neurotree: A collaborative, graphical database of the academic genealogy of neuroscience. PloS one, 7(10).

DiCarlo, J.J., Zoccolan, D., and Rust, N.C. (2012). How does the brain solve visual object recognition? Neuron 73, 415–434.

Eisenreich, B. R., Akaishi, R., & Hayden, B. Y. (2017). Control without controllers: toward a distributed neuroscience of executive control. Journal of cognitive neuroscience, 29(10), 1684–1698.

Farashahi, S., Azab, H., Hayden, B., & Soltani, A. (2018). On the flexibility of basic risk attitudes in monkeys. Journal of Neuroscience, 2260–17.

Farashahi, S., Donahue, C. H., Hayden, B. Y., Lee, D., & Soltani, A. (2019). Flexible combination of reward information across primates. Nature human behaviour, 3(11), 1215–1224.

Fellows, L. K. (2006). Deciding how to decide: ventromedial frontal lobe damage affects information acquisition in multi-attribute decision making. Brain, 129(4), 944–952.

Gelman, A., Stern, H. S., Carlin, J. B., Dunson, D. B., Vehtari, A., & Rubin, D. B. (2013). Bayesian data analysis. Chapman and Hall/CRC.

Hare, T. A., Schultz, W., Camerer, C. F., O’Doherty, J. P., & Rangel, A. (2011). Transformation of stimulus value signals into motor commands during simple choice. Proceedings of the National Academy of Sciences, 201109322.

Hayden, B. Y. (2019). Why has evolution not selected for perfect self-control?. Philosophical Transactions of the Royal Society B, 374(1766), 20180139.

Hayden, B., & Gallant, J. (2013). Working memory and decision processes in visual area v4. Frontiers in neuroscience, 7, 18.

Hayden, B. Y., & Platt, M. L. (2010). Neurons in anterior cingulate cortex multiplex information about reward and action. Journal of Neuroscience, 30(9), 3339–3346.

Heilbronner, S. R., & Hayden, B. Y. (2016). Dorsal anterior cingulate cortex: a bottom-up view. Annual review of neuroscience, 39, 149–170.

Heilbronner, S. R., & Hayden, B. Y. (2016). The description-experience gap in risky choice in nonhuman primates. Psychonomic bulletin & review, 23(2), 593–600.

Hayden, B. Y., & Moreno-Bote, R. (2018). A neuronal theory of sequential economic choice. Brain and Neuroscience Advances, 2, 2398212818766675.

Hunt, L. T., Behrens, T. E., Hosokawa, T., Wallis, J. D., & Kennerley, S. W. (2015). Capturing the temporal evolution of choice across prefrontal cortex. Elife, 4, e11945.

Hunt, L. T., Dolan, R. J., & Behrens, T. E. (2014). Hierarchical competitions subserving multi-attribute choice. Nature neuroscience, 17(11), 1613.

Hunt, L. T., & Hayden, B. Y. (2017). A distributed, hierarchical and recurrent framework for reward-based choice. Nature Reviews Neuroscience, 18(3), 172.

Hunt, L. T., Kolling, N., Soltani, A., Woolrich, M. W., Rushworth, M. F., & Behrens, T. E. (2012). Mechanisms underlying cortical activity during value-guided choice. Nature neuroscience, 15(3), 470.

Hunt, L. T., Malalasekera, W. N., de Berker, A. O., Miranda, B., Farmer, S. F., Behrens, T. E., & Kennerley, S. W. (2018). Triple dissociation of attention and decision computations across prefrontal cortex. Nature neuroscience, 21(10), 1471.

Hosokawa, T., Kennerley, S. W., Sloan, J., & Wallis, J. D. (2013). Single-neuron mechanisms underlying cost-benefit analysis in frontal cortex. Journal of Neuroscience, 33(44), 17385–17397.

Kable, J. W., & Glimcher, P. W. (2009). The neurobiology of decision: consensus and controversy. Neuron, 63(6), 733–745.

Kahnt, T., Heinzle, J., Park, S. Q., & Haynes, J. D. (2011). Decoding different roles for vmPFC and dlPFC in multi-attribute decision making. Neuroimage, 56(2), 709–715.

Kennerley, S. W., Dahmubed, A. F., Lara, A. H., & Wallis, J. D. (2009). Neurons in the frontal lobe encode the value of multiple decision variables. Journal of cognitive neuroscience, 21(6), 1162–1178.

Kim, S., Hwang, J., & Lee, D. (2008). Prefrontal coding of temporally discounted values during intertemporal choice. Neuron, 59(1), 161–172.

Klein-Flügge, M. C., & Bestmann, S. (2012). Time-dependent changes in human corticospinal excitability reveal value-based competition for action during decision processing. Journal of neuroscience, 32(24), 8373–8382.

Kriegeskorte, N., Mur, M., & Bandettini, P. A. (2008). Representational similarity analysis-connecting the branches of systems neuroscience. Frontiers in systems neuroscience, 2, 4.

Levy, D. J., & Glimcher, P. W. (2012). The root of all value: a neural common currency for choice. Current opinion in neurobiology, 22(6), 1027–1038.

Louie, K., Grattan, L. E., & Glimcher, P. W. (2011). Reward value-based gain control: divisive normalization in parietal cortex. Journal of Neuroscience, 31(29), 10627–10639.

Maxwell, S. E., & Delaney, H. D. (1993). Bivariate median splits and spurious statistical significance. Psychological bulletin, 113(1), 181.

Montague, P. R., & Berns, G. S. (2002). Neural economics and the biological substrates of valuation. Neuron, 36(2), 265–284.

Noguchi, T., & Stewart, N. (2018). Multialternative decision by sampling: A model of decision making constrained by process data. Psychological review, 125(4), 512.

O’Doherty, J. P. (2014). The problem with value. Neuroscience & Biobehavioral Reviews, 43, 259–268.

O’Neill, M., & Schultz, W. (2018). Predictive coding of the statistical parameters of uncertain rewards by orbitofrontal neurons. Behavioural brain research.

Padoa-Schioppa, C. (2011). Neurobiology of economic choice: a good-based model. Annual review of neuroscience, 34, 333–359.

Padoa-Schioppa, C., & Assad, J. A. (2006). Neurons in the orbitofrontal cortex encode economic value. Nature, 441(7090), 223.

Passino, K. M., Seeley, T. D., & Visscher, P. K. (2008). Swarm cognition in honey bees. Behavioral Ecology and Sociobiology, 62(3), 401–414.

Paxinos, G., Huang, X. F., & Toga, A. W. (2000). The rhesus monkey brain in stereotaxic coordinates.

Pirrone, A., Azab, H., Hayden, B. Y., Stafford, T., & Marshall, J. A. (2018). Evidence for the speed–value trade-off: Human and monkey decision making is magnitude sensitive. Decision, 5(2), 129.

Raghuraman, A. P., & Padoa-Schioppa, C. (2014). Integration of multiple determinants in the neuronal computation of economic values. Journal of Neuroscience, 34(35), 11583–11603.

Rangel, A., Camerer, C., & Montague, P. R. (2008). A framework for studying the neurobiology of value-based decision making. Nature reviews neuroscience, 9(7), 545.

Rangel, A., & Hare, T. (2010). Neural computations associated with goal-directed choice. Current opinion in neurobiology, 20(2), 262–270.

Roesch, M. R., & Olson, C. R. (2004). Neuronal activity related to reward value and motivation in primate frontal cortex. Science, 304(5668), 307–310.

Roesch, M. R., & Olson, C. R. (2003). Impact of expected reward on neuronal activity in prefrontal cortex, frontal and supplementary eye fields and premotor cortex. Journal of Neurophysiology, 90(3), 1766–1789.

Rushworth, M. F., Noonan, M. P., Boorman, E. D., Walton, M. E., & Behrens, T. E. (2011). Frontal cortex and reward-guided learning and decision-making. Neuron, 70(6), 1054–1069.

Rustichini, A., & Padoa-Schioppa, C. (2015). A neuro-computational model of economic decisions. Journal of neurophysiology, 114(3), 1382–1398.

Schall, J. D., Stuphorn, V., & Brown, J. W. (2002). Monitoring and control of action by the frontal lobes. Neuron, 36(2), 309–322.

Seeley, T. D. (2009). The wisdom of the hive: the social physiology of honey bee colonies. Harvard University Press.

Seeley, T. D., & Buhrman, S. C. (1999). Group decision making in swarms of honey bees. Behavioral Ecology and Sociobiology, 45(1), 19–31.

Seeley, T. D., Visscher, P. K., Schlegel, T., Hogan, P. M., Franks, N. R., & Marshall, J. A. (2012). Stop signals provide cross inhibition in collective decision-making by honeybee swarms. Science, 335(6064), 108–111.

Shizgal, P. (1997). Neural basis of utility estimation. Current opinion in neurobiology, 7(2), 198–208.

Sleezer, B. J., Castagno, M. D., & Hayden, B. Y. (2016). Rule encoding in orbitofrontal cortex and striatum guides selection. Journal of Neuroscience, 36(44), 11223–11237.

Sleezer, B. J., LoConte, G. A., Castagno, M. D., & Hayden, B. Y. (2017). Neuronal responses support a role for orbitofrontal cortex in cognitive set reconfiguration. European Journal of Neuroscience, 45(7), 940–951.

Sleezer, B. J., & Hayden, B. Y. (2016). Differential contributions of ventral and dorsal striatum to early and late phases of cognitive set reconfiguration. Journal of cognitive neuroscience, 28(12), 1849–1864.

So, N. Y., & Stuphorn, V. (2010). Supplementary eye field encodes option and action value for saccades with variable reward. Journal of Neurophysiology, 104(5), 2634–2653.

Stewart, N., Chater, N., & Brown, G. D. (2006). Decision by sampling. Cognitive psychology, 53(1), 1–26.

Strait, C. E., Blanchard, T. C., & Hayden, B. Y. (2014). Reward value comparison via mutual inhibition in ventromedial prefrontal cortex. Neuron, 82(6), 1357–1366.

Strait, C. E., Sleezer, B. J., Blanchard, T. C., Azab, H., Castagno, M. D., & Hayden, B. Y. (2015). Neuronal selectivity for spatial positions of offers and choices in five reward regions. Journal of neurophysiology, 115(3), 1098–1111.

Strait, C. E., Sleezer, B. J., & Hayden, B. Y. (2015). Signatures of value comparison in ventral striatum neurons. PLoS biology, 13(6), e1002173.

Thura, D., & Cisek, P. (2016). Modulation of premotor and primary motor cortical activity during volitional adjustments of speed-accuracy trade-offs. Journal of Neuroscience, 36(3), 938–956.

Tversky, A. (1969). Intransitivity of preferences. Psychological review, 76(1), 31.

Tversky, A., & Kahneman, D. (1974). Judgment under uncertainty: Heuristics and biases. science, 185(4157), 1124–1131.

Wang, M. Z., & Hayden, B. Y. (2017). Reactivation of associative structure specific outcome responses during prospective evaluation in reward-based choices. Nature communications, 8, 15821.

Wunderlich, K., Rangel, A., & O’Doherty, J. P. (2009). Neural computations underlying action-based decision making in the human brain. Proceedings of the National Academy of Sciences, 106(40), 17199–17204.

Yoo, S. B. M., & Hayden, B. Y. (2018). Economic choice as an untangling of options into actions. Neuron, 99(3), 434–447.

Yoo, S. B. M., Tu, J. C., Piantadosi, S. T., & Hayden, B. Y. (2020). The neural basis of predictive pursuit. Nature Neuroscience, 1–8.

Yoo, S. B. M., & Hayden, B. Y. (2020). The Transition from Evaluation to Selection Involves Neural Subspace Reorganization in Core Reward Regions. Neuron, 105(4), 712–724.

